# Brief Report: Prognostic relevance of 3q amplification in squamous cell carcinoma of the lung

**DOI:** 10.1101/2022.09.28.509728

**Authors:** Fawzi Abu Rous, Pin Li, Shannon Carskadon, Sunny RK Singh, Rebecca Chacko, Hassan Abushukair, Shirish Gadgeel, Nallasivam Palanisamy

## Abstract

**Introduction:** Amplification of 3q is the most common genetic alteration identified in squamous cell carcinoma of the lung (LUSC) with the most frequent amplified region being 3q26-3q28.

**Methods:** In this analysis, we aim to describe the prognostic relevance of 3q by focusing on a minimal common region (MCR) of amplification within 3q constituted by 25 genes. We analyzed 511 cases of LUSC from The Cancer Genome Atlas (TCGA) and included 476 in the final analysis.

**Results:** We identified a 25-gene MCR that was amplified in 221 (44.3%) cases and was associated with better disease specific-survival (DSS) (NR versus 9.25 years; 95% CI [5.24-NR]; log-rank p=0.011) and a progression-free interval (PFI) of 8 years (95% CI [5.1-NR]) versus 4.9 years (95% [3.5-NR]) (Log-rank p=0.020). Multivariable analysis revealed MCR amplification was associated with improved DSS and PFI.

**Conclusion:** Amplification of the 25-gene MCR within 3q was present in 44% of this cohort mainly composed of Caucasian patients with early-stage LUSC. This analysis is a strong indicator of the prognostic relevance of the 25-gene MCR within 3q. We are further evaluating its prognostic relevance in a racially diverse patient population with advanced LUSC.

## Introduction

Lung cancer is the second most common cancer and the leading cause of cancer-related deaths worldwide. The most common type of lung cancer is adenocarcinoma (LUAD) followed by squamous cell carcinoma (LUSC), both of which comprise a majority of non-small cell lung cancer (NSCLC)^1^. The outcomes of patients with lung cancer have changed significantly since the identification of driver mutations and the introduction of targeted therapies and immune checkpoint inhibitors (ICI)^2^. Driver mutations that can be therapeutically targeted are more commonly detected in LUAD, and rarely detected in patients with LUSC^3^. Patients with advanced LUSC are typically treated with platinum doublet chemotherapy and ICIs without the benefit of targeted therapies; therefore, survival of LUSC patients is inferior to those with LUAD^2^. Biomarker studies are urgently needed in LUSC for prognostic and predictive purposes.

Amplification of 3q is the most common genetic alteration observed in LUSC with the most frequently amplified region being 3q26-3q28^4^. It has been described in preinvasive and invasive LUSC and represents one of the most striking differences between LUSC and LUAD^5^. Interestingly, 3q amplification is prevalent and carries prognostic relevance in head and neck, cervical, and esophageal squamous cell cancers^6^. The distal amplified area of 3q includes genes that are key in squamous differentiation such as SOX-2, p63, and PIK3CA which explains the prevalence and importance of this genetic alteration in squamous cell carcinomas^6^. Many studies reported the significance of 3q amplification in LUSC and investigated the entire 3q region (∼40 genes) or focused on 1-2 genes of known biological relevance across tumor types such as SOX-2, p63, PI3KCA, FGFR1^7-10^. In our study, we have identified a minimal common region (MCR) of amplification constituted of 25 genes within 3q using The Cancer Genome Atlas (TCGA) LUSC dataset and examined its prognostic relevance.

## Methods

We conducted analysis of 511 LUSC cases in TCGA and identified amplification of MCR (chr3:181,711,924-183,428,101) ∼1.7MB within a large, amplified region between (chr3:170,169,718-187,736,569) ∼17.5MB on chromosome 3q (Figure 1). MCR was estimated based on the smallest genomic interval amplified in most of the cases. MCR constitutes of 25 genes that were found to be frequently amplified and share a common downstream PIK3CA pathway that was found to be altered in 71% of the cases (Figure 2). The MCR contains the following 25 genes: *SOX2, ATP11B, MCCC1, LAMP3, MCF2L2, B3GNT5, KLHL6, KLHL24, YEATS2, PARL, MAP6D1, ABCC5, HTR3D, HTR3E, EIF2B5, DVL3, AP2M1, ABCF3, VWA5B2, ALG3, CAMK2N2, ECE2, PSMD2, EIF4G1, FAM131A*.

**Figure 1:**
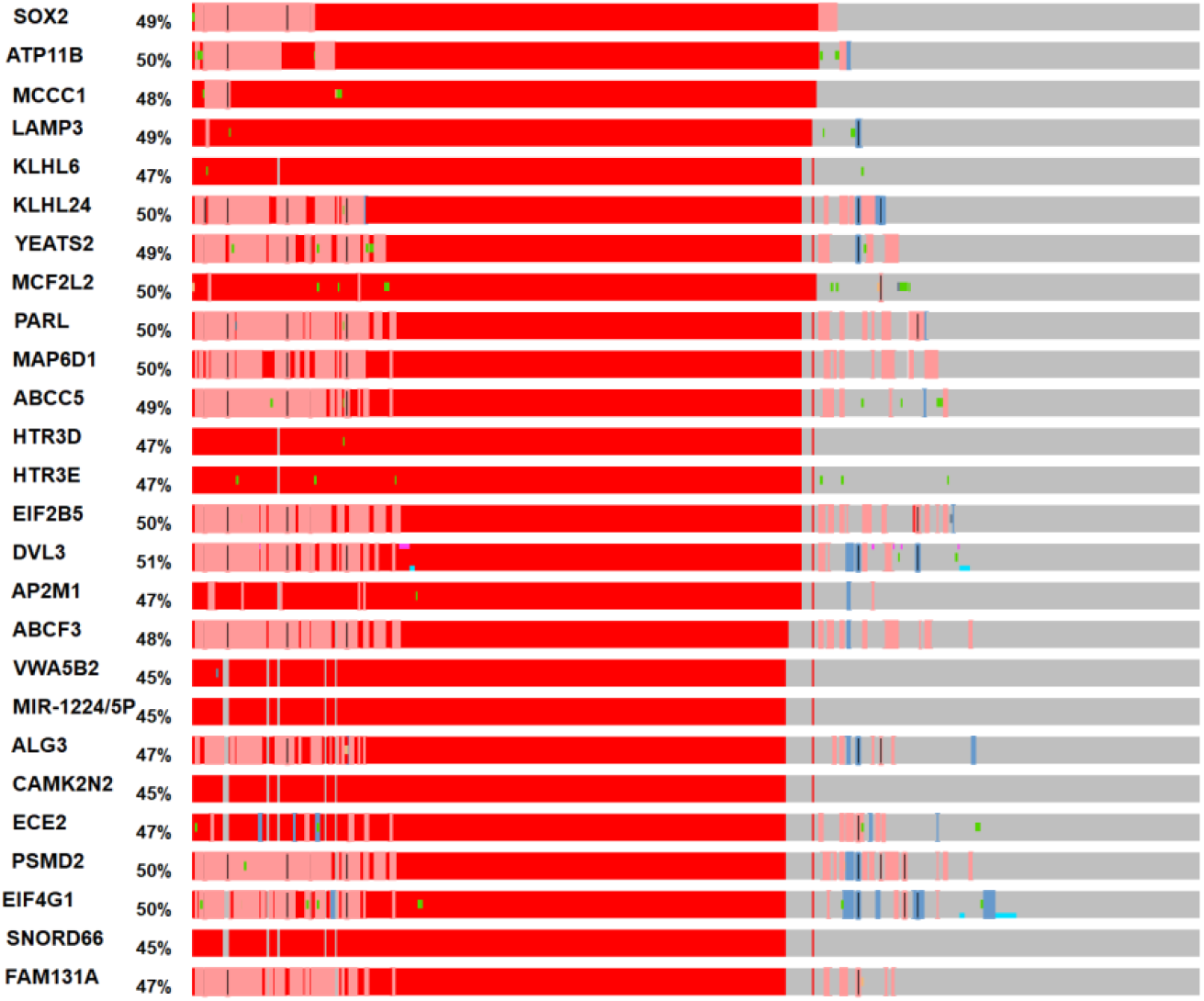
Minimal Common Region of Amplification, analyzed 17.5MB region on 3q (chr3:170,169,718-187,736,569) between PHC3 and BCL6 MCR was identified as 1.7MB region (chr3:181,711,924-183,428,101) from SOX2 to MCF2L2

**Figure 2:**
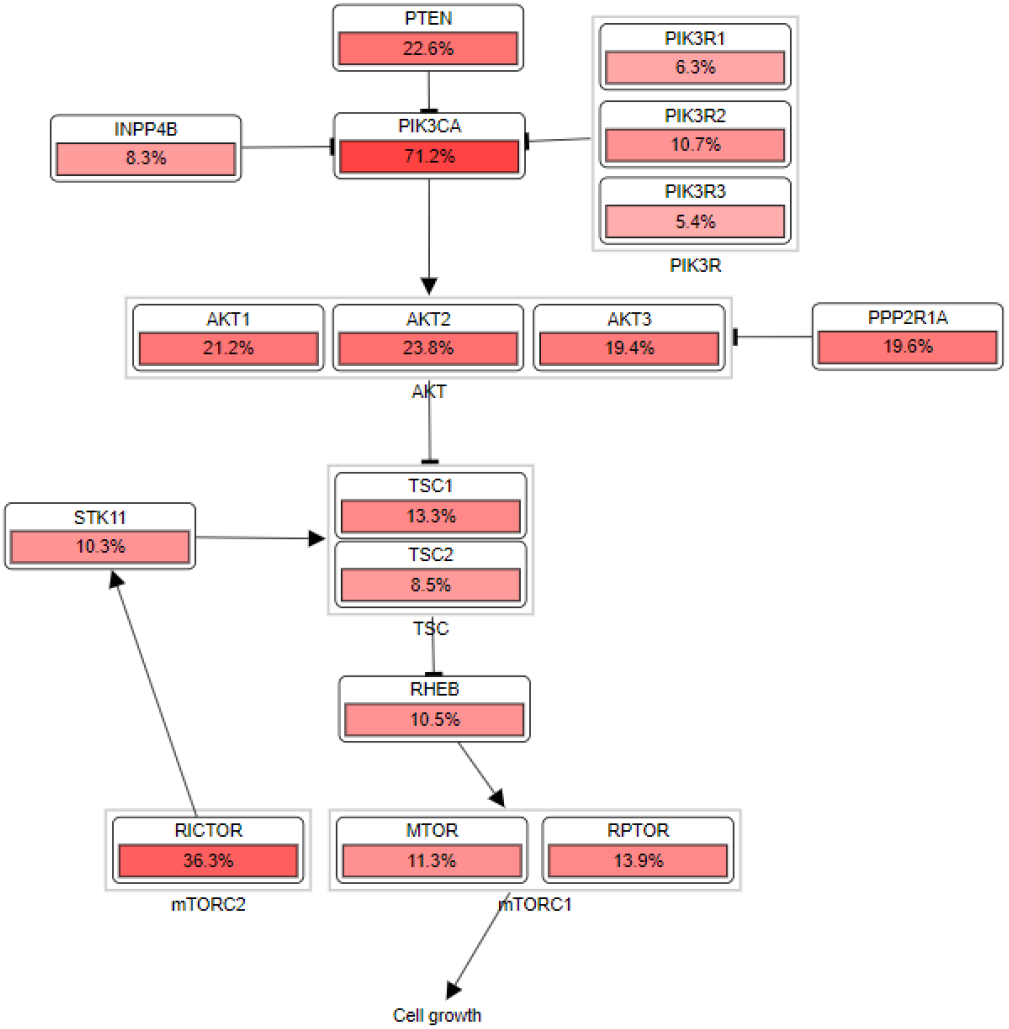
PI3K is a common pathway shared by 25 genes within MCR

### Patients

We identified 511 patients with LUSC in TCGA and excluded 5 patients with synchronous disease. Outcome data was available for 499 patients and extracted from Clinical Data Resources. We downloaded gene copy number variation data (amplification/ deletion) for the 25 genes in MCR within 3q and CDKN2A from TCGA Firehose Legacy set in CBioPortal. We excluded 3 patients with no molecular profiling, and 20 patients with partial MCR amplification. The final analysis was done using clinical and molecular data for 476 patients.

## Statistical Analysis

Patient clinical characteristics were summarized with continuous variable by mean and standard deviation and categorical variable by count and percentage. Comparison by MCR amplification was conducted by Fisher’s exact test. DSS (disease-specific survival) is the date of initial diagnosis until the date of death from the disease. The progression free interval (PFI) is the time from diagnosis until the first progression (recurrence of disease or death with disease) or the time last known alive without progression. DSS and PFI was compared by MCR amplification status using Kaplan-Meier method and log-rank test. Multivariable Cox-proportional hazard (Cox-PH) models were used to evaluate the association of DSS and PFI with age, gender, stage, smoking, CDKN2A deletion and MCR amplification. All analysis were conducted using R statistical software version 4.0.2. Two tailed p-value<0.05 was used for statistical significance.

## Results

### Patient characteristics

In the overall cohort, the mean age was 67.4 years, 73.9% were males and 26.1% were females. Race information was available for 367 cases, 332 (69.7%) identified as white, 26 (5.5%) identified as African American (AA), and 9 (1.9%) identified as Asian. Only 6 patients (1.3%) had stage IV disease, whereas 48.5% had stage I, 33.1% had stage II, and 17.2% had stage III. Regarding smoking status, 26.9% were current smokers, 67.4% were former smokers and 3.4% were never smokers (Table 1).

**Table 1:** Patient characteristics for the overall cohort as well as MCR-amplified and non-amplified patients

| Characteristics | Overall | MCR amplified | MCR non-amplified |
| --- | --- | --- | --- |
| Total N (%) | 476 | 221 (46.4) | 255 (53.6) |
| Age at diagnosis – Mean (SD) | 67.4 (8.6) | 66.6 (8.0) | 68.1 (9.1) |
| Gender – N (%) |  |  |  |
| Female | 124 (26.1) | 58 (26.2) | 66 (25.9) |
| Male | 352 (73.9) | 163 (73.8) | 189 (74.1) |
| Race – N (%) |  |  |  |
| White | 332 (69.7) | 154 (69.7) | 178 (69.8) |
| African American | 26 (5.5) | 12 (5.4) | 14 (5.5) |
| Asian | 9 (1.9) | 3 (1.4) | 6 (2.4) |
| Unknown | 109 (22.9) | 52 (23.5) | 57 (22.4) |
| Stage – N (%) |  |  |  |
| I | 229 (48.5) | 109 (49.8) | 120 (47.4) |
| II | 156 (33.1) | 74 (33.8) | 82 (32.4) |
| III | 81 (17.2) | 34 (15.5) | 47 (18.6) |
| IV | 6 (1.3) | 2 (0.9) | 4 (1.6) |
| Smoking – N (%) |  |  |  |
| Current | 128 (26.9) | 62 (28.1) | 66 (25.9) |
| Former | 321 (67.4) | 150 (67.9) | 171 (67.1) |
| Never | 16 (3.4) | 4 (1.8) | 12 (4.7) |
| Unknown | 11 (2.3) | 5 (2.3) | 6 (2.4) |
| CDKN2A Status – N (%) |  |  |  |
| No alteration | 342 (72.2) | 147 (66.8) | 195 (76.8) |
| Homozygous Deletion | 132 (27.8) | 73 (33.2) | 59 (23.2) |
| TP53 Status – N (%) |  |  |  |
| No alteration | 319 (67) | 180 (70.6) | 139 (62.9) |
| Alteration | 157 (33) | 75 (29.4) | 82 (37.1) |
| PIK3CA Status – N (%) |  |  |  |
| No amplification | 217 (45.6) | 215 (97.3) | 2 (0.8) |
| Amplification | 259 (54.4) | 6 (2.7) | 253 (99.2) |

### MCR Amplification

MCR is constituted of 25 genes that were frequently amplified and signal through PI3K downstream pathway which was altered in 71% of the cases (Figure 2). Full 25-gene MCR amplification was found in 221 (44.3%) patients whereas 255 (51%) patients had no amplification. MCR amplification was associated with more CDKN2A homozygous deletion which was detected in 70 (31.7%) with MCR amplification and 56 (22%) without amplification (OR: 1.64, p=0.021; 95% CI: 1.07-2.54). TP53 alteration was less detected in MCR-amplified cases, 75 (29.4%) vs 82 (37.1%) (OR: 0.707, p=0.079; 95% CI 0.473-1.056). MCR amplification was strongly associated with PIK3CA amplification which was detected in 215 (97.3%) of MCR-amplified cases compared to 2 (0.8%) of non-amplified cases (OR: 4469, p<0.001) which is most likely due to the proximity of PIK3CA (located on 3q26.32) with MCR (starts from 3q26.33).

### Outcomes

The Median DSS of MCR-amplified cases was significantly longer than non-amplified cases (NR versus 9.25 years; 95% CI [5.24-NR]; log-rank p=0.011). Median PFI for amplified cases was 8 years (95% CI [5.1-NR]) versus 4.9 years (95% [3.5-NR]) for non-amplified cases (Log-rank p=0.020) (Figure 3). In multivariable analysis, stage III was associated with worse DSS (HR 3.10, 95% CI [1.80-5.33], p<0.001) and PFI (HR 2.37 95% CI [1.53-3.66], p<0.001), and MCR amplification was associated with improved DSS (HR 0.59, 95% CI [0.38-0.94], p=0.025) and PFI (HR 0.67 95% CI [0.47-0.95], p=0.024). Compared to current smokers, former smokers had a lower risk of death (HR 0.60, 95% CI [0.37-0.97], p=0.035) and progression (HR 0.67, 95% CI [0.46-0.98], p=0.039) (Table 2).

**Table 2:** Multivariable analysis of disease specific survival (DSS) and progression free interval (PFI)

| Variable |  | DSS HR | PFI HR |
| --- | --- | --- | --- |
| Age | 67.4 (mean) | 0.96 (0.75-1.23, p=0.741) | 1.04 (0.86-1.26, p=0.695) |
| Gender | Female | REF | REF |
|  | Male | 1.27 (0.75-2.16, p=0.367) | 1.08 (0.73-1.60, p=0.714) |
| Stage | I | REF | REF |
|  | II | 1.62 (0.96-2.74, p=0.072) | **1.63 (1.10-2.40, p=0.015)** |
|  | III/IV | **3.10 (1.80-5.33, p<0.001)** | **2.37 (1.53-3.66, p<0.001)** |
| CDKN2A | No Alteration | REF | REF |
|  | Homozygous Deletion | 1.11 (0.68-1.84, p=0.674) | 1.16 (0.78-1.71, p=0.467) |
| 3q AMP | No | REF | REF |
|  | **AMP** | **0.59 (0.38-0.94, p=0.025)** | **0.67 (0.47-0.95, p=0.024)** |
| Smoking | Current | REF | REF |
|  | Former | **0.60 (0.37-0.97, p=0.035)** | **0.67 (0.46-0.98, p=0.039)** |
|  | Never | 1.69 (0.49-5.89, p=0.409) | 1.89 (0.78-4.59, p=0.159) |
|  | Unknown | 0.62 (0.19-2.06, p=0.432) | 1.17 (0.50-2.78, p=0.716) |

**Figure 3:**
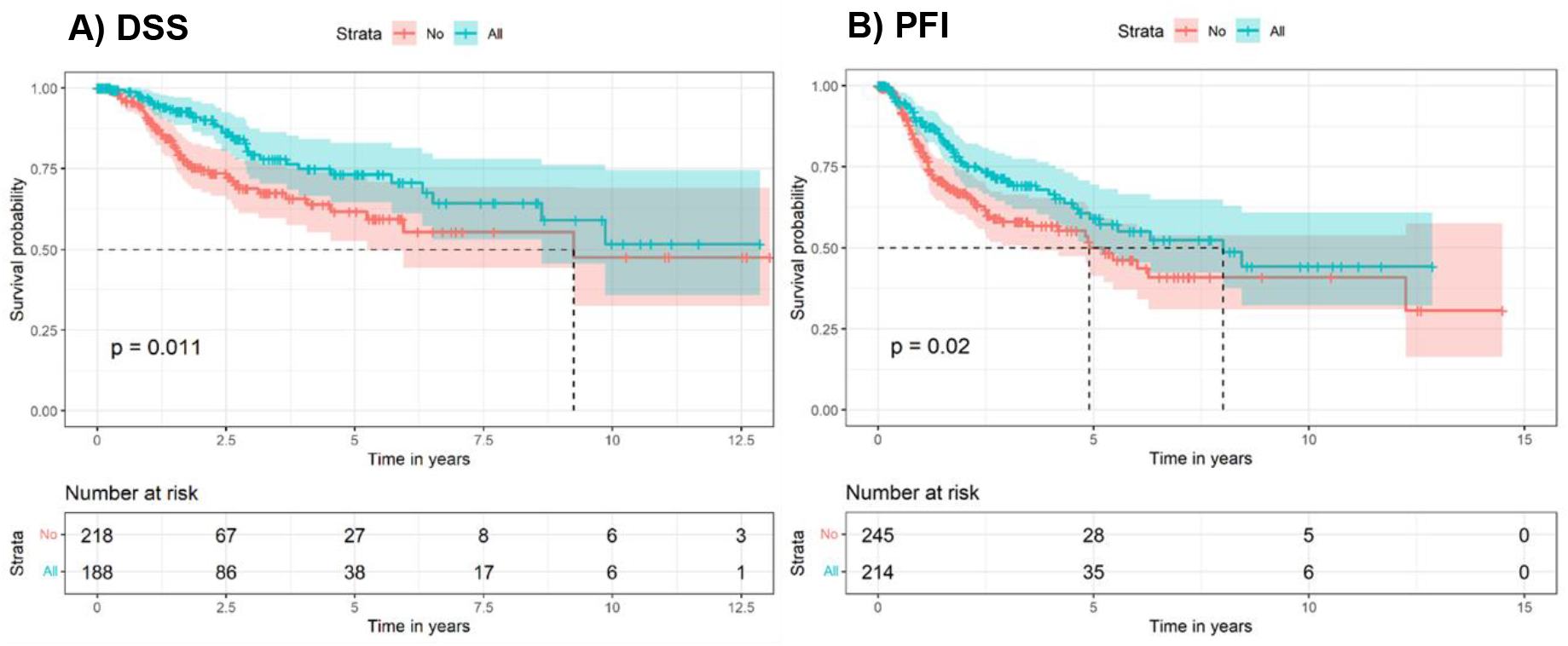
A) Disease-specific Survival, B) Progression Free Interval for MCR-amplified and non-amplified patients.

## Discussion

Unlike LUAD, patients with LUSC are rarely found to have single-driver genetic alterations that can be therapeutically targeted. In one study, 13% of LUSC samples were found to have at least one potentially targetable genetic alteration compared to 66% of LUAD samples^11^. Smoking as a risk factor plays a major role in the diverse genomic landscape of LUSC, making the task of identifying a single-driver genetic alteration very challenging. LUSC is characterized by significant genomic complexity and a high rate of mutations (8.1 mutations per megabase), as shown by a comprehensive genomic analysis of LUSC by TCGA. This analysis showed recurrent mutations in 11 genes, including *TP53*, in almost all analyzed tumors. It also reported selective 3q amplification as the most common genetic alteration in LUSC and it is the most notable difference between LUSC and LUAD^12^.

Many genes in this region have been evaluated as potential therapeutic targets and their role as biomarkers were investigated as well. *SOX-2* is a “lineage-survival oncogene” that promotes squamous identity and encodes a transcription factor that has an important role in cellular proliferation and expansion. SOX-2 was reported to be altered in 17.7% of the samples in one cohort, and its overexpression was associated with a better median overall survival (68 versus 35 months, P=0.036)^7^.

*P63* is another gene within the 3q amplicon encoding a transcription factor that leads to cellular apoptosis after activating *TP53* genes. The *P63* gene encodes 6 splicing variants, ΔNp63 being the most common. ΔNp63 lacks the NH2-terminal domain altering its function to promote growth rather than apoptosis. Patients with tumors exhibiting genomic amplification of p63 had significantly prolonged survival compared to P63 non-amplified patients^8^. PI3CKA amplification, which promotes cancer growth and migration, was detected in 37% of LUSC and was reported to be associated with worse survival in a Japanese cohort of 92 patients^13^. FGFR1 is a known oncogene of the FGFR family, it was found to be amplified in 16% of the samples in 226 patients; however, no association with survival was detected^10^. FXR1 was identified as a potential driver in the 3q amplicon by Qian et al, they also reported that FXR1 overexpression is a poor prognostic factor in multiple solid malignancies including NSCLC^14^.

It has been proposed that multiple genes in this area of amplification may have a synergistic effect on the progression of LUSC. Qian et al. identified a 12-gene signature within 3q from LUSC dataset in TCGA. They reported a negative correlation between their 12-gene signature and gene groups involved in immune checkpoints and immune-related processes indicating a suppressed immune pathway in their patient population^15^.

Evaluating a single gene within the 3q region has led to contradicting results, as described above. Hence, we evaluated the region as a whole and identified MCR within 3q where all 25 genes were frequently amplified. We found this region to be amplified in 44.3% of LUSC in a cohort that was composed mainly of Caucasians (90.2%) with early-stage disease I-III (98.5%) and lacked adequate representation of the minority patient population and patients with advanced lung cancer. MCR amplification was associated with better DSS and PFI. In MCR-amplified cases, all 25 genes within MCR were amplified but not necessarily highly expressed at the mRNA level. We identified 4 genes that were amplified and highly expressed: ABCC5, AP2M1, EIF4G1, and PSMD2. EIF4G1 has been shown by our group to be a poor prognostic marker and a potential therapeutic target in multiple cancers. ABCC5 is involved in recurrent gene rearrangement in LUSC, based on our observations from the TCGA database, which requires further validation and characterization. The function of these amplified/ highly expressed genes will be investigated in future studies.

This approach has been used by Qian et al as described above; however, the 25-gene MCR not only carries a prognostic value but also a potential therapeutic significance as PI3K is the most altered downstream pathway among MCR-amplified cases. PI3K pathway is a known target and can potentially offer new therapeutic options for patients with LUSC, PI3Kinase inhibitors are currently used in breast cancer patients and certain hematologic malignancies. These agents have been evaluated in NSCLC patients with limited efficacy. There is a need to conduct a focused assessment of these agents in LUSC patients with MCR amplification.

## Conclusion

Amplification on 25-gene MCR within 3q is a good prognostic marker in a patient population with early-stage LUSC. It also offers opportunities for targeted therapeutics given the common PI3K pathway between MCR genes and the high prevalence of PI3KCA amplification in MCR-amplified cases. We are conducting a validation retrospective study to examine the prognostic and predictive value of 3q MCR amplification using tissue samples from a racially diverse patient population with advanced LUSC using a FISH probe created at Palanisamy’s lab to detect MCR amplification (Figure 4).

**Figure 4:**
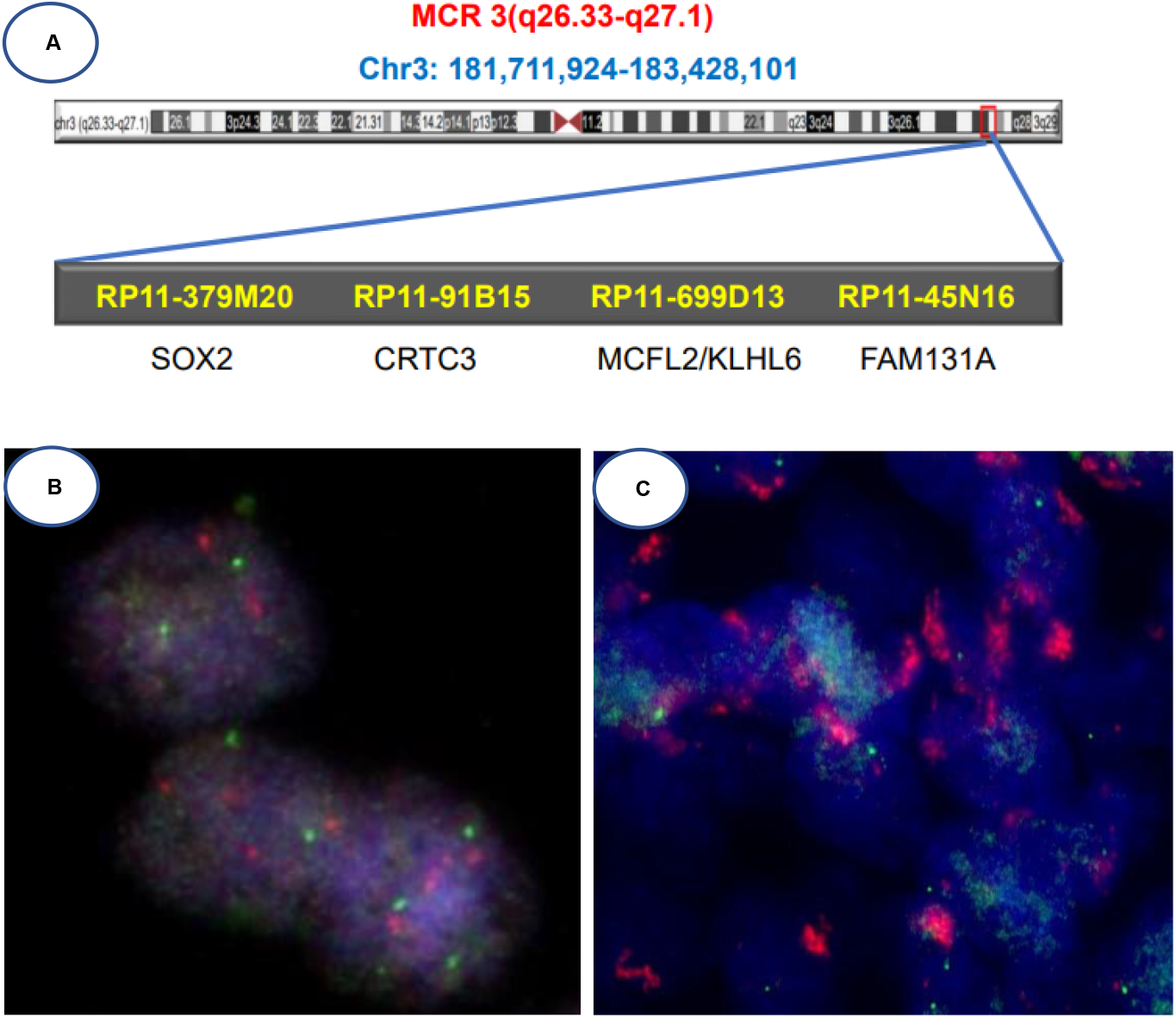
A) 3q MCR FISH probe, example of FISH results in tissue samples: B) non-amplified, C) amplified

## Acknowledgments

The authors thank Laila Possion, PhD (Department of Public Health Sciences, Henry Ford Health) for providing assistance with statistical analysis.

